# Alternative oxidase1a and 1d limit proline-dependent oxidative stress and aid salinity recovery in Arabidopsis

**DOI:** 10.1101/2021.08.02.454800

**Authors:** Glenda Guek Khim Oh, Brendan M. O’Leary, Santiago Signorelli, A. Harvey Millar

## Abstract

A link between Pro catabolism and mitochondrial reactive oxygen species production has been established across eukaryotes and in plants increases in leaf respiration rates have been reported following Pro exposure. Here we investigated how alternative oxidases (AOXs) of the mitochondrial electron transport chain accommodate the large, atypical flux resulting from Pro catabolism and limit oxidative stress during Pro breakdown in mature Arabidopsis leaves. Following Pro treatment, AOX1a and AOX1d accumulate at transcript and protein levels, with AOX1d approaching the level of the typically dominant AOX1a isoform. We therefore sought to determine the function of both AOX isoforms under Pro respiring conditions. Oxygen consumption rate measurements in *aox1a* and *aox1d* leaves suggested these AOXs can functionally compensate for each other to establish enhanced AOX catalytic capacity in response to Pro. Generation of *aox1a.aox1d* lines showed complete loss of AOX proteins and activity upon Pro treatment, yet full respiratory induction in response to Pro remained possible via the cytochrome pathway. However, *aox1a.aox1d* leaves suffered increased levels of oxidative stress and damage during Pro metabolism compared to WT or the single mutants. During recovery from salt stress, when high rates of Pro catabolism occur naturally, photosynthetic rates in *aox1a.aox1d* recovered slower than WT or the single *aox* lines, showing that both AOX1a and AOX1d are beneficial for cellular metabolism during Pro drawdown following osmotic stress. This work provides physiological evidence of a beneficial role for *AOX1a* but also the less studied *AOX1d* isoform in allowing safe catabolism of alternative respiratory substrates like Pro.

**One sentence summary:** The alternative oxidase of plant mitochondria contributes to Pro catabolism by preventing oxidative stress in the electron transport chain and this aids recovery of leaf metabolic rates following salinity stress.

## INTRODUCTION

Plant respiration is a tightly regulated metabolic network that is capable of adapting to changing cellular needs and growth conditions by altering its metabolic behaviour (Plaxton and Podestá, 2006, O’Leary et al., 2019a). Through the complete or partial oxidation of substrates, plant respiration serves three main purposes: ATP synthesis, biosynthetic precursor synthesis and cellular redox balancing (O’Leary et al., 2019a). The principal respiratory substrates in non-photosynthesizing plant tissues are carbohydrates, metabolised sequentially via glycolysis and the tricarboxylic acid (TCA) cycle, yielding their reducing power to feed the mitochondrial electron transport chain (mETC) (O’Leary and Plaxton, 2016). However, under certain conditions, often associated with starvation and stress, plants rely on alternative respiratory substrates such as amino acids or derivatives of fatty acids, which are metabolised through various additional respiratory pathways leading to the mETC (Millar et al., 2011, Rasmusson and Møller, 2011, Cabassa-Hourton et al., 2016, Launay et al., 2019, O’Leary et al., 2019b). In photosynthesizing cells, mitochondrial respiration also has a clear function to enhance photosynthesis by participating in photorespiration (glycine oxidation) and oxidising excess reductant exported by the plastid (Raghavendra and Padmasree, 2003, Tcherkez et al., 2017, Del-Saz et al., 2018). Because of its size and functional complexity, our understanding of the control and regulation of plant respiration as a network is limited (Sweetlove et al., 2010). In turn, our ability to understand how variability in respiratory metabolism contributes to plant performance in nature (O’Leary et al., 2017) or in the field (Amthor et al., 2019, Reynolds et al., 2021) is constrained.

Like other aspects of the respiratory network, the plant mETC is highly flexible from a metabolic standpoint (Rasmusson et al., 2008, Millar et al., 2011). Firstly, multiple non-proton pumping alternative enzyme exist which bypass components of the standard mETC (*i.e*., complexes I-IV) (Millar et al., 2011). In particular, plant mitochondria contain various isoforms of alternative oxidase (AOX), an enzyme that catalyses the transfer of electrons directly from ubiquinol to O_2_, thereby bypassing complexes III and IV of the standard mETC and partly uncoupling respiration from ATP generation (Millar et al., 2011, Vanlerberghe, 2013). Because AOX activity appears to constitute an energetically wasteful process, researchers have long sought to understand the countervailing benefits of AOX activity in non-thermogenic tissues. Electron transport through the mETC contributes to ROS production, especially when electron flux is high or the activity of mETC components is disrupted (Møller, 2001, Huang et al., 2016). Under these scenarios, AOX activity could function to reduce the redox poise of the ubiquinone pool and other mETC components to thereby to mitigate ROS formation (Millar et al., 2001). Previous reports have shown that the overexpression of AOX in transgenic tobacco cells (Maxwell et al., 1999) and drought-treated tobacco leaves (Dahal and Vanlerberghe, 2017), activation of AOX in isolated durum wheat mitochondria (Pastore et al., 2001) and upregulation of AOX in salt-stressed Arabidopsis seedlings (Gong et al., 2020) all resulted in less ROS abundance and oxidative damage. A second potential benefit of AOX activity is that it allows the mETC to run faster and be more responsive when redox balancing, albeit with lower rates of ATP production. AOX also supports photosynthetic activity and it would appear to do this by allowing the mETC to rapidly dissipate excess reducing power, thus contributing to overall cellular redox balance (Vishwakarma et al., 2015, Jiang et al., 2019). Despite some remaining uncertainty about the precise benefits of AOX expression and function, available evidence suggests its presence is favourable for plant growth (Vanlerberghe, 2013, Del-Saz et al., 2018, Selinski et al., 2018b).

In Arabidopsis, the AOX gene family consists of five genes: *AOX1a* – *AOX1d* and *AOX2*. Analysis of the distribution and conservation AOX family genes throughout the plant kingdom is complicated, but *AOX1d* genes, *AOX1a-c* genes and *AOX2* genes represent three separate clades that are present in many plant families (Costa et al., 2014). Transcriptomic and proteomic analyses have consistently indicated that AOX1a is the dominant AOX isoform in Arabidopsis in all vegetative tissues (Clifton et al., 2006). *AOX1b* is expressed during early flowering stages while *AOX1c* and *AOX2* are expressed at low levels regardless of developmental stages (Clifton et al., 2006). Both *AOX1a* and *AOX1d* demonstrate a clear pattern of elevated expression under a range stress conditions such as salinity (Van Aken et al., 2009, Feng et al., 2013) and low nitrogen stress (Watanabe et al., 2010) and also in response to chemical inhibition of mETC complexes, such as with antimycin A (Strodtkotter et al., 2009). Manipulation of *AOX1a* expression in Arabidopsis under diverse environmental treatments such as cold (Fiorani et al., 2005), drought (Giraud et al., 2008), high CO_2_ concentrations (Gandin et al., 2012), high arsenic (Demircan et al., 2020) and ecologically relevant doses of UV-B (Garmash et al., 2020) have highlighted the potential of *AOX1a* to enable acclimation to unfavourable growth conditions. *AOX1a* expression is particularly strongly upregulated by chemical disruption of complex II and III, and has been linked to mitochondrial retrograde signalling (Sweetlove et al., 2002, Clifton et al., 2005, Schwarzländer et al., 2011, Umbach et al., 2012) and the alleviation of excess reducing power arising from impediments to normal mETC activity (Jiang et al., 2019). Comparable genetic studies with altered expression of AOX2 have been performed on *Glycine max* (Chai et al., 2012) and *Cerasus humilis* (Sun et al., 2021). Selinski et al. (2018a) showed that Arabidopsis AOX isoforms are differentially activated by various TCA cycle intermediates, suggesting sub-functionalisation of the AOX gene family. Still there is limited information on the physiological function of AOX genes other than *AOX1a* in plants (Clifton et al., 2006, Strodtkotter et al., 2009, Konert et al., 2015).

Changes in cellular metabolite levels strongly influence respiratory rates in leaves (O’Leary et al., 2017, O’Leary et al., 2019b). In particular, external Pro exposure is followed by cellular accumulation, which induces a strong stimulation of Pro respiratory catabolism in Arabidopsis leaves, leading to a doubling of respiration rate over 14 hours (O’Leary et al., 2019b). Pro is an atypical respiratory substrate in plants whose accumulation and metabolism under osmotic stress and recovery is well-established (Yoshiba et al., 1995, Hare et al., 1998, Verbruggen and Hermans, 2008, Verslues and Sharma, 2010). Here, we suspected that Pro’s ability to potently stimulate respiration presented an opportunity to study metabolic flexibility at the level of the mETC. The two-step pathway of Pro oxidative catabolism to Glu is understood. The mitochondrial enzyme proline dehydrogenase (ProDH), oxidizes Pro to pyrroline-5-carboxylate (P5C) and likely supplies electrons directly to ubiquinol via its FADH cofactor (Szabados and Savouré, 2010, Servet et al., 2012, Cabassa-Hourton et al., 2016, Launay et al., 2019). In the second step, P5C spontaneously transforms into glutamate-5-semialdehyde (GSA), which is oxidised to Glu by mitochondrial P5C dehydrogenase (P5CDH) with the concomitant reduction of NADH (Forlani et al., 1997, Verslues and Sharma, 2010, Trovato et al., 2019). Pro catabolism increases ROS production *via* ProDH activity In mammalian cells (Donald et al., 2001) and *C. elegans* (Zarse et al., 2012), while in plants, the absence of P5CDH led to increased mitochondrial ROS following Pro exposure (Miller et al., 2009). What is currently unclear is how mETC components would adjust to accommodate the rapid and non-standard electron flux change resulting from scenarios requiring large scale Pro catabolism. In this study, we provide evidence that Pro exposure specifically upregulates AOX1d and AOX1a expression, greatly increasing AOX capacity within the mETC. Our results indicate that *AOX1a* and *AOX1d* both act to limit oxidative stress during Pro catabolism and thereby facilitate plant recovery from osmotic stress.

## RESULTS

### Leaf exposure to Pro leads to an increase in AOX1a and AOX1d transcripts and protein abundances

Pro treatment leads to a pronounced *PDH*-dependent stimulation of leaf night time oxygen consumption rates (OCR) (Fig 1A) (O’Leary et al., 2019b), a situation that may evoke strong deviations from typical energy and redox balances. We hypothesised that increased leaf OCR following Pro exposure requires stimulation of mechanisms that uncouple the mETC from ATP production and retain redox balance (*e.g*., AOX or uncoupling protein (UCP) expression). We used multiple reaction monitoring (MRM) to quantify a variety of mitochondrial protein abundances in WT control and Pro-exposed leaf samples, including 24 components of the classical electron transport chain, and 43 components of TCA cycle and enzymes of organic and amino acid metabolism (Supplemental Table 1). Eight proteins showed significant abundance differences in response to Pro, with four being upregulated and four being down regulated. AOX1d and glutamate dehydrogenase 2 (GDH2) increased four-fold, AOX1a increased two-fold, and uncoupling protein 1 (UCP1) displayed a minor induction (Fig 1B). On the other hand, isocitrate dehydrogenase subunit 1, pyruvate dehydrogenase complex subunit 1-2 and E1 α subunit, and succinyl CoA ligase β subunit showed modest but significant decreases after Pro treatment (Supplemental Table 1). Notably, GDH can function downstream of Pro catabolism and catalyses glutamate deamination to 2-OG, thus GDH2 likely functions to connect Pro catabolism to the TCA cycle. Analysis of the abundance of 24 components distributed across the Complexes I-V of the classical ETC showed no Pro-dependent changes in abundance (Supplemental Table 1).

**Figure 1:**
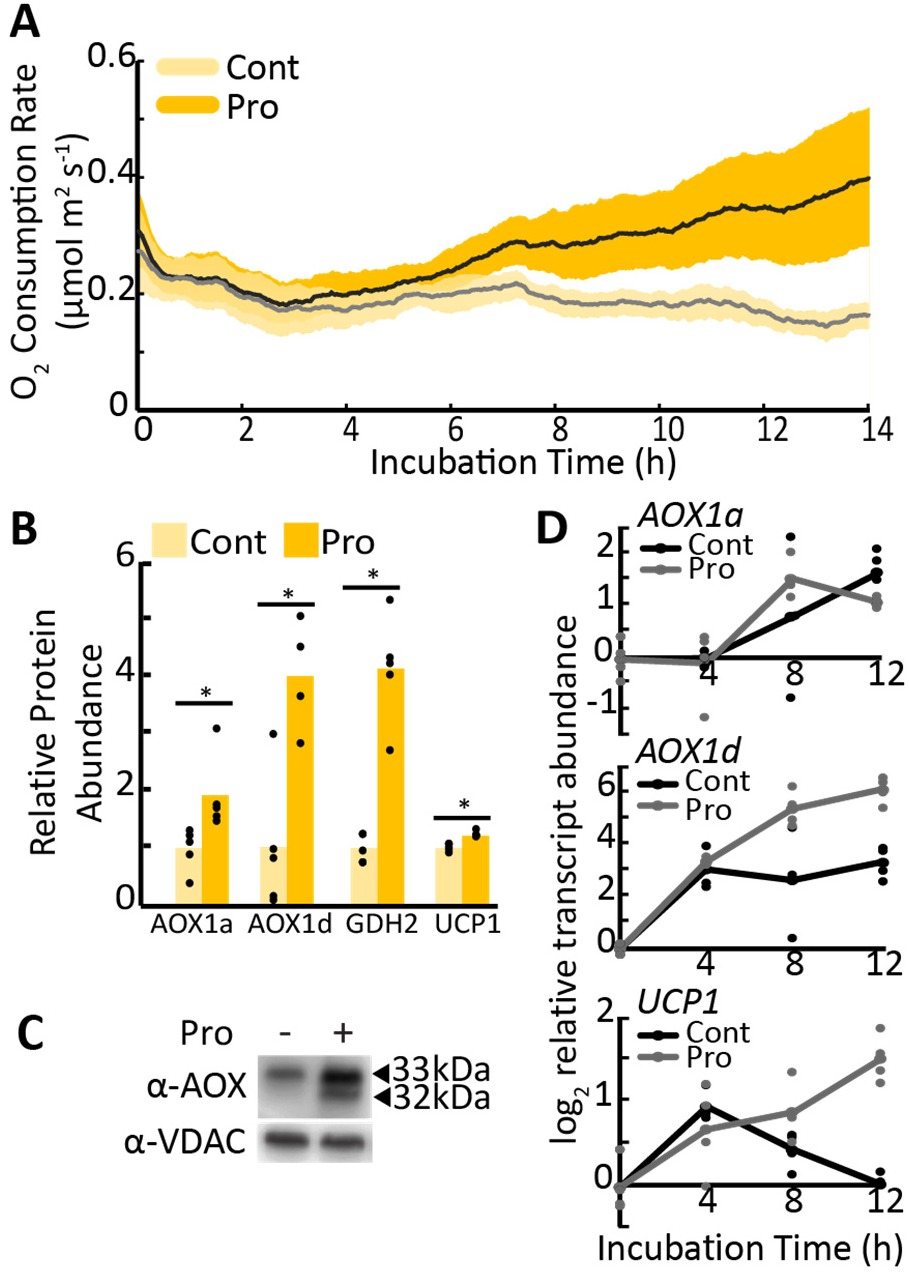
Transcript and protein abundance of AOX1a and AOX1d after Pro exposure in leaf tissues. **(A)** Time-dependent OCR stimulation in single leaf discs floated on buffer solution with or without Pro. Shaded area indicated the 95% confidence intervals (n=6). **(B)** Mitochondrial proteins that significantly increased in abundance following Pro treatment of leaf discs, shown relative to non-treated samples. Asterisks represent significant differences between treatments (*t*-test, *n* ≥ 4; p < 0.05). **(C)** Immunoblot analysis of enriched mitochondrial fractions from WT leaf tissue incubated in the presence or absence of Pro. Bands corresponding to AOX isoforms are indicated. **(D)** Transcript analysis of *AOX1a, AOX1d* and *UCP1* in control and Pro-treated leaf discs. Lines represent mean values; replicate data points are shown (*n* ≥ 3 independent biological replicates).

The large upregulation of multiple AOX isoforms was confirmed by immunoblotting of enriched mitochondrial samples from Pro treated leaf discs (Fig 1C). Transcript analysis of leaf discs also confirmed the time-dependent upregulation of *AOX1a, AOX1d* and *UCP1* in response to Pro treatment, suggesting the enhanced expression of these uncoupling proteins was controlled at least partly at the level of transcription (Fig 1D). By contrast, *AOX1b, AOX1c* and *AOX2* transcripts did not increase in response to Pro (Supplemental Fig 1). Together these data indicated that alterations to AOX expression represented the largest change to mETC functionality we could find upon Pro exposure.

We distinguished two AOX immunoreactive bands in WT leaves that increased in abundance following Pro exposure (Fig 1C). Similar doublet bands have been previously observed on AOX immunoblots of isolated Arabidopsis mitochondria following various treatments, and Konert et al. (2015) identified an upper band as AOX1a and a lower band as AOX1d using tandem mass-spectrometry. To establish the molecular nature of the two AOX bands in our treatments, we probed enriched mitochondrial samples isolated from an established *aox1a* line (SAIL_303_D04) and two *aox1d* lines, *aox1d 1* (SALK_203986) and *aox1d 2* (GABI_529D11) that we characterised here (Fig 2A and Supplemental Fig 2A). The immunoblot banding pattern observed in the *aox1a* and *aox1d* samples confirmed the association of the upper and lower immunoreactive bands with AOX1a (33 kDa) and AOX1d (32 kDa), respectively (Fig 2B). Both *aox1d* lines lacked any detectable AOX1d transcript and protein, and AOX1d expression was only detectable upon Pro treatment in WT and *aox1a* plants (Fig 2B - D). Although *AOX1d* transcript levels remained much lower than those of *AOX1a*, the AOX1d protein accumulated to appreciable amounts under Pro treatment. Therefore, the *aox1a* line was not an effective AOX knockout in this instance. To achieve an effective AOX knockout, two independent *aox1a.aox1d* lines were constructed. *AOX1a* and *AOX1d* transcripts were absent (Fig 2D and Supplemental Fig 2A) and no AOX protein could be detected by immunoblotting of enriched mitochondrial samples following leaf Pro treatment in both *aox1a.aox1d* lines (Fig 2B – C & Supplemental Fig 2B). There was also no observable growth phenotype between WT and any of the *aox* lines under our standard growth conditions (Supplemental Fig 2C). We repeated the mitochondrial MRM analysis on samples from *aox1a.aox1d 1* leaves and confirmed the loss of AOX1a and AOX1d abundance (Fig 2C, Supplemental Table 1). *aox1a.aox1d* plants displayed a minor induction of *UCP1* transcript in both control and Pro-exposed leaf tissues indicating a potential compensation for the lack of AOX.

**Figure 2:**
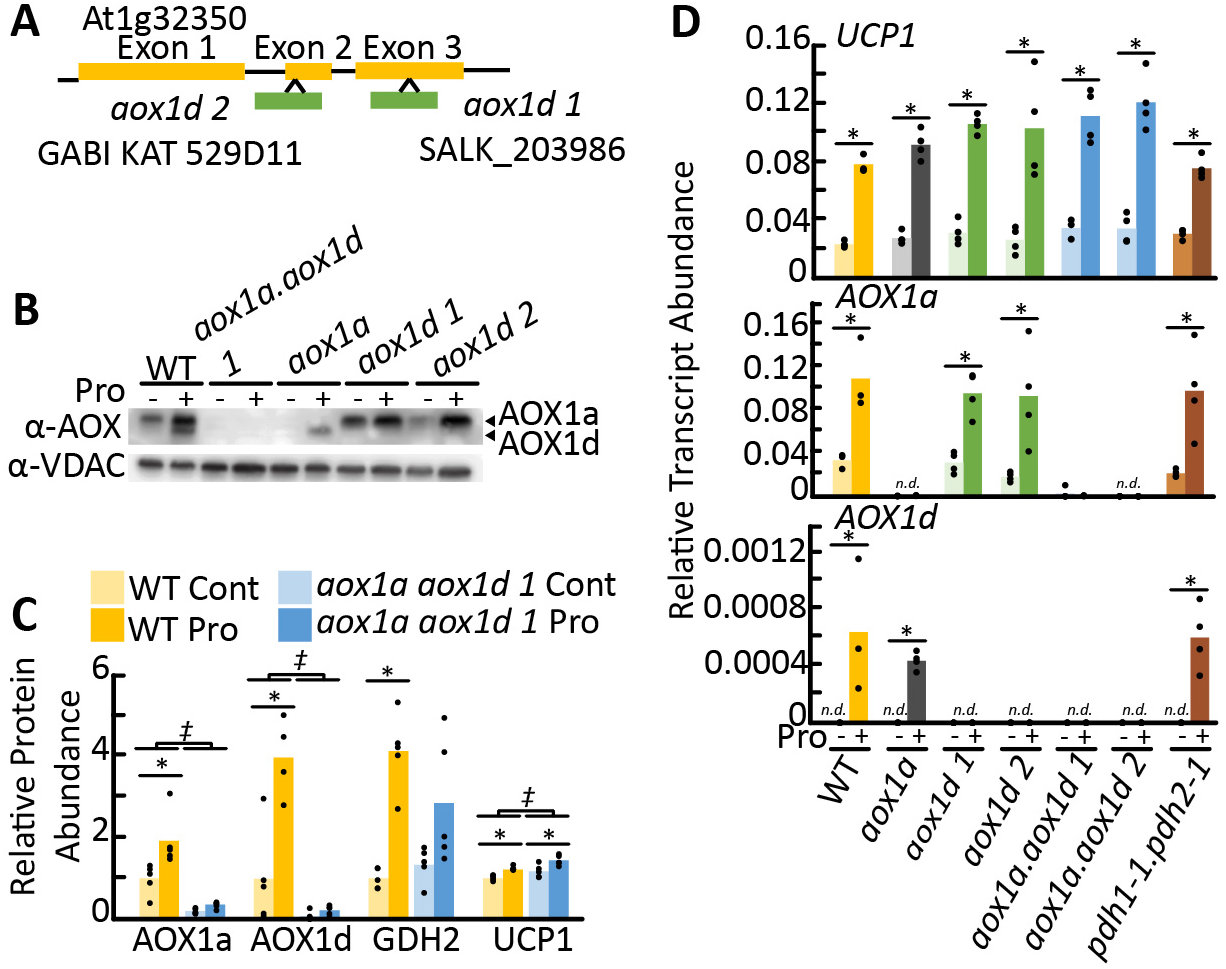
Characterisation of *aox1a, aox1d* and *aox1a.aox1d* lines. **(A)** Gene model of AOX1d (At1g32350) indicating T-DNA insertion locations corresponding to SALK_203986 and GABI KAT_529D11. Genotyping data are show in Supplemental Figure 2. (**B**) Immunoblot analysis of enriched mitochondrial fractions from leaf tissue pre-incubated in the presence or absence of Pro. Each lane contains 20 μg of protein. Bands corresponding to AOX1a and AOX1d are indicated. **(C)** Multiple reaction monitoring protein abundance measurements. Relative abundance is calculated in comparison to wild-type non-treated samples. Asterisks represent significant differences between treatments, double daggers represent significant differences between genotypes (two-way ANOVA, *n* ≥ 4; p < 0.05). (**D**) Transcript analysis of *AOX1a, AOX1d* and *UCP1* following 14 h Pro treatment of leaf discs. Bars represent average transcript abundance; replicate data points are shown (*n* ≥ 3). Asterisks represent significant differences between treatments (ANOVA; p < 0.05).

We investigated whether the presence of Pro itself or the metabolism of Pro via the mETC acted as the signal to stimulate AOX expression. Accumulation of *AOX1a, AOX1d* and *UCP1* transcripts following Pro treatment was unaffected in *pdh1-1.pdh2-1* leaves that lack expression of ProDH (Fig 2D). Previous work using these mutants has demonstrated that ProDH is needed for Pro catabolism via the mETC (Cabassa-Hourton et al., 2016, O’Leary et al., 2019b). This result suggests that elevated Pro level even in the absence of high Pro respiratory flux is sufficient to signal AOX expression.

### Pro-dependent respiratory flux is maintained in the absence of AOX

To understand if either AOX1a or AOX1d had a role in Pro-dependent respiration, we measured OCR in WT, *aox1a, aox1d and aox1a.aox1d* leaf discs exposed to Pro. Similar Pro-inductions of OCR were observed in all genotypes (Fig 3A-B). This indicates that AOX is not required, and that the cytochrome pathway (COX) capacity is sufficient to carry electron transport chain flux resulting from high rates of Pro catabolism.

**Figure 3:**
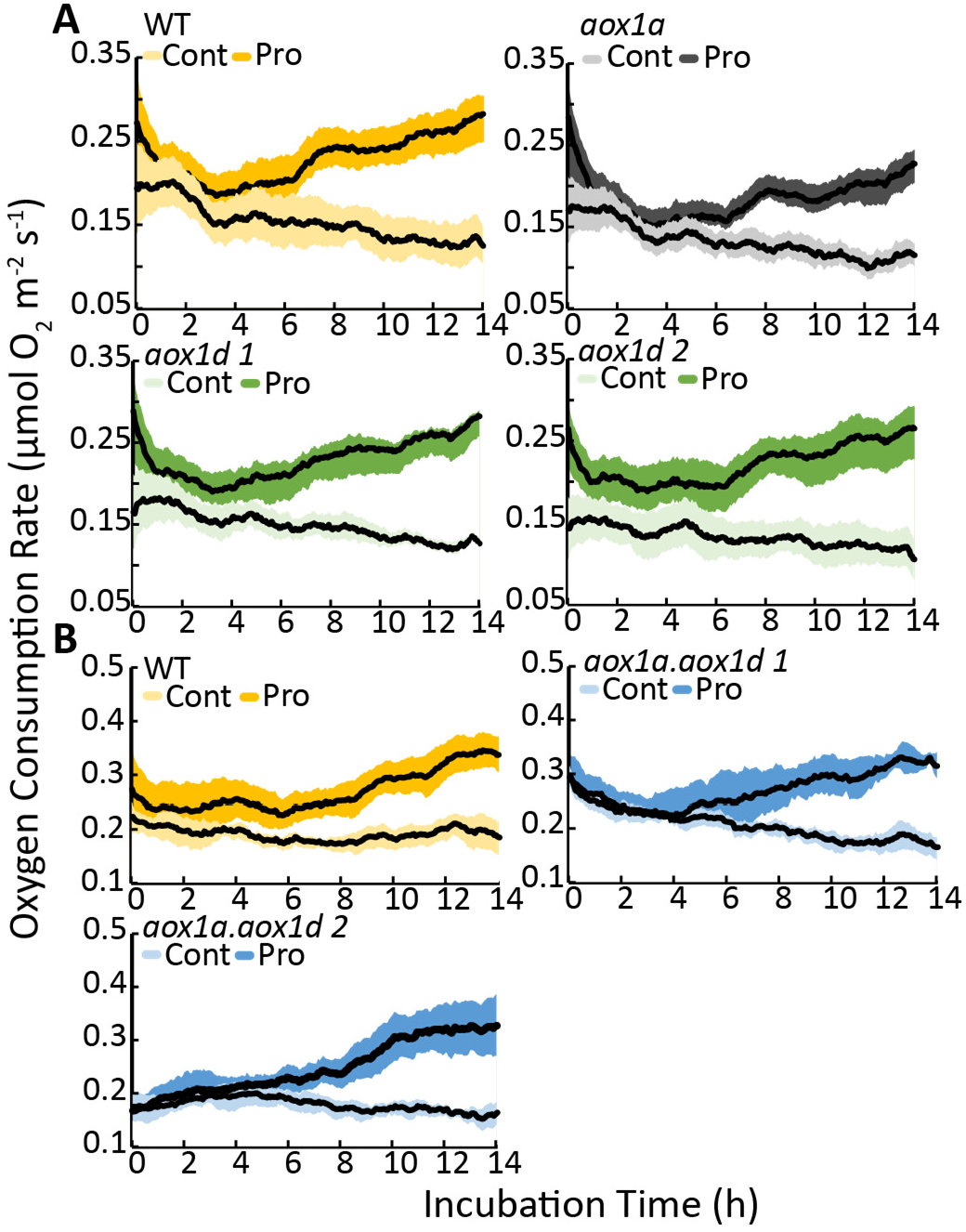
The effect of the loss of AOX1a or AOX1d on Pro stimulated R_N_ in AOX knockout lines. Time-dependent stimulation in respiration after Pro exposure. Single leaf discs of (**A**) WT, *aox1a, aox1d 1, aox1d 2* and **(B)** WT, *aox1a.aox1d 1* and *aox1a.aox1d 2* were floated on 10 mM Pro respiration buffer. Control treated leaf discs were floated on respiration buffer absent of Pro. Traces show moving average respiration rates (*n* = 6). The shaded area shows the 95% confidence intervals.

We would note that additional experiments conducted with commonly used AOX inhibitors, SHAM and nPG, produced contradictory results. While the presence of SHAM did not affect Pro uptake (Supplemental Fig 3A), SHAM completely inhibited Pro-dependent respiratory stimulation in leaf discs of all genotypes including both *aox1a.aox1d* lines (Supplemental Fig 3B). This effect of SHAM was therefore independent of AOX expression level in leaf tissue. Furthermore, OCR of enriched mitochondrial fractions derived from control and Pro-treated leaf tissue was not substantially inhibited by nPG when either Pro or NADH and succinate were available as respiratory substrates (Supplemental Fig 3C). As SHAM can affect multiple oxidases beyond AOX (Kahn and Zakin, 2000), we hypothesize that additional SHAM sensitive oxidases are involved in Pro signalling in leaf tissue. Therefore, the use of genetics rather than AOX inhibitors is preferred here to allow for a more direct interpretation of Pro-dependent respiratory fluxes over longer time periods.

### AOX1d contributes to increased alternative pathway capacity upon Pro treatment

We wanted to verify whether AOX isoforms and AOX1d in particular, could substantially contribute to mETC capacity in leaves following Pro exposure. OCR measurements of Pro pre-treated leaf discs were performed using Clark-type oxygen electrodes to observe rapid OCR changes, wherein KCN additions were used to measure AOX capacity. In agreement with above Q2 experiments, there was an approximately 2-fold stimulation of pre-KCN OCR resulting from Pro pre-treatment and Pro stimulation of OCR was not significantly different in the absence of AOX1a and/or AOX1d (Fig 4A-C).

**Figure 4:**
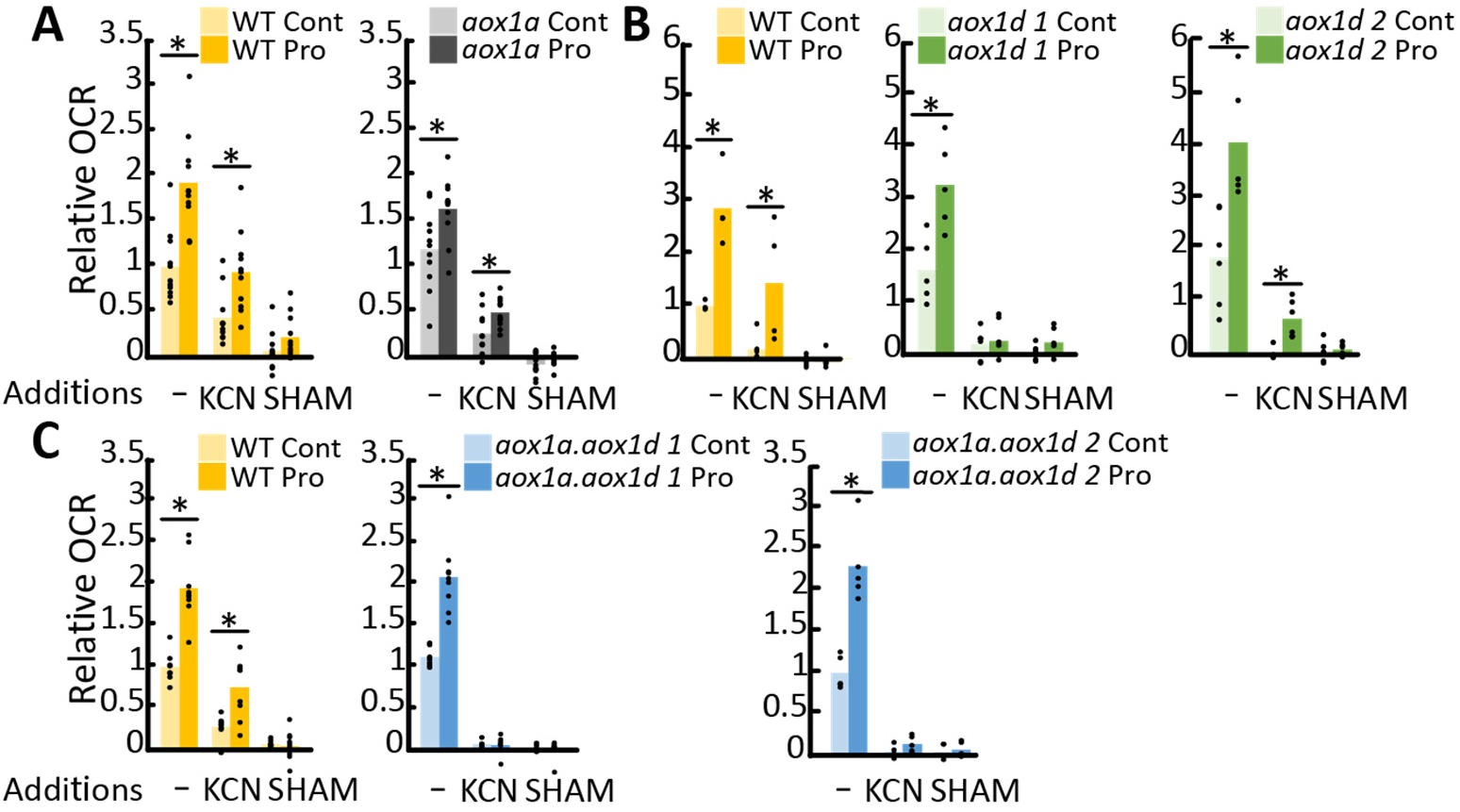
Both AOX1a and AOX1d contribute to the increase in AOX capacity after Pro exposure. OCR measurements of control or Pro-treated leaf discs were made before (-) and after the sequential addition of KCN and SHAM. Separate experiments compared **(A)** *aox1a*, **(B)** *aox1d* lines and **(C)** *aox1a.aox1d* lines to WT. Data are expressed relative to uninhibited OCR in corresponding WT control samples. Replicate data points are shown (*n* ≥ 4 independent biological replicates) Asterisks represent significant differences between treatments (*t*-test; p < 0.05).

The addition of KCN revealed a substantial induction of AOX capacity by Pro-pre-treatment in WT leaf discs. By comparison, AOX capacity (following KCN addition) was significantly less in *aox1a* (Fig 4A) and *aox1d* (Fig 4B), and absent in both *aox1a.aox1d* lines (Fig 4C). In all instances, KCN resistant respiration was rapidly inhibited by SHAM. These results indicated that AOX capacity is upregulated in response to Pro due to contributions from both AOX1a and AOX1d.

### AOX1a and AOX1d are required to limit Pro-induced oxidative stress

To understand the potential benefit of enhanced AOX capacity on respiratory metabolism following Pro exposure, we performed a time course metabolomics analyses in *aox1a.aox1d* and WT leaf discs following Pro treatments for 0, 4 or 8 h (Fig 5 and Supplemental Table 2). Pro treatment caused a strong accumulation of both ascorbate (Asc; reduced) and dehydroascorbate (DHA; oxidised ascorbate), indicating an activation of antioxidant mechanisms. Notably, *aox1a.aox1d* samples displayed significantly less Asc across all treatments indicating possible differences in oxidative stress (Fig 5 and Supplemental Table 2). The absence of AOX also led to minor significant decreases in citrate, fumarate, 2-oxoglutarate, isocitrate, valine, glucose and pyruvate across all treatment types (Fig 5 and Supplemental Table 2).

**Figure 5:**
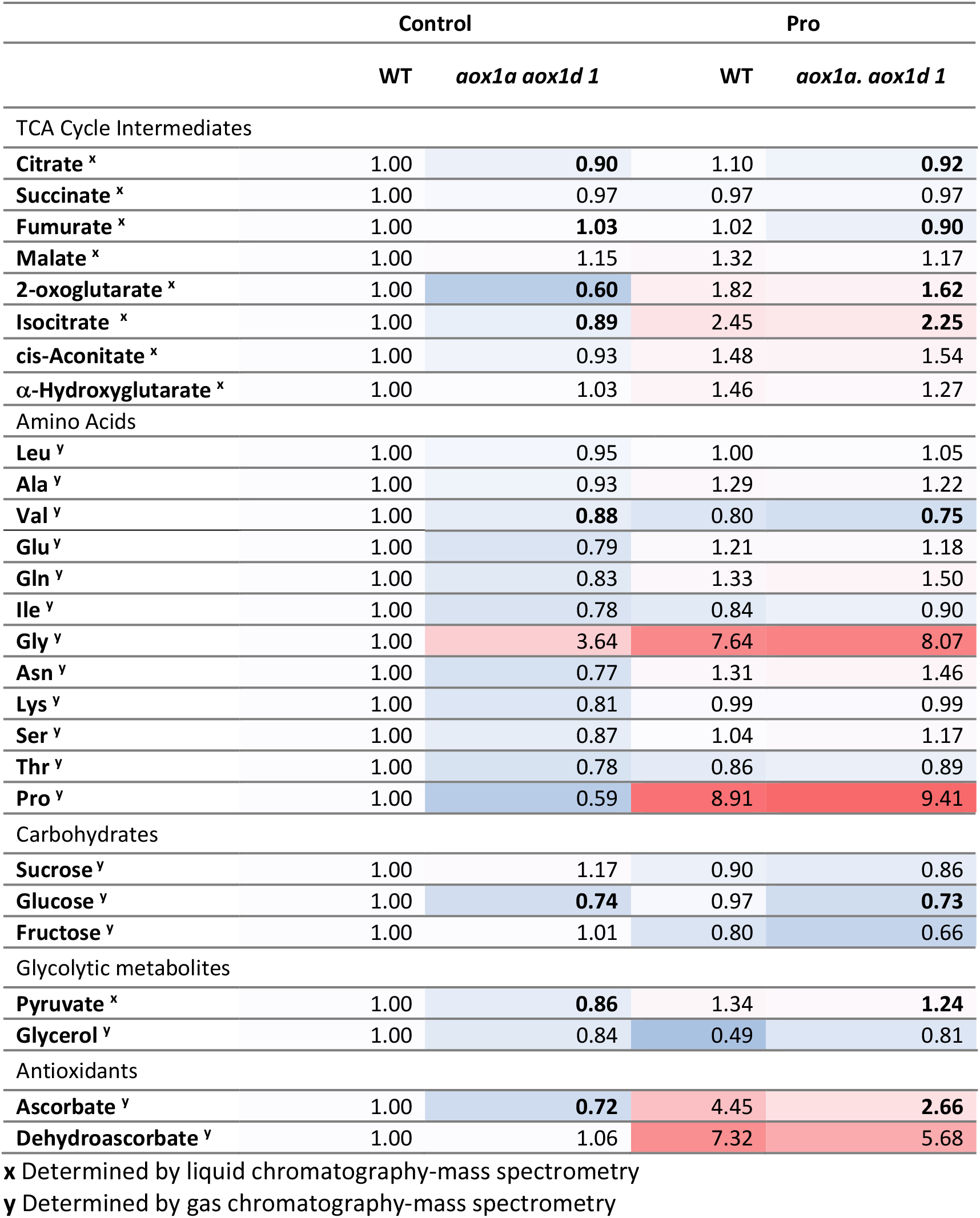
Quantification of AOX- and Pro-dependent changes in cellular metabolites. Measurement of cellular metabolites in WT and *aox1a.aox1d 1* leaf discs following 4 h Pro treatment. Average metabolite abundance was calculated relative to WT control samples. Red and blue boxes refer to higher and lower metabolite abundances. Boldface indicates significant differences between WT and *aox1a.aox1d* samples (three-way ANOVA, p<0.05). See supplemental table 1 for full dataset.

Quantification of actual Asc:DHA ratios using GC-MS is not possible as the oxidation state of Asc is not strictly preserved and the two metabolites are measured without absolute quantification. Therefore, we conducted additional measures to verify the redox status of Asc pools and oxidative damage in Pro treated leaf discs. Following 4 h of Pro exposure, Asc redox status in *aox1a.aox1d* lines, but not single *aox1a* or *aox1d* knockouts, was more oxidised in comparison to WT (Fig 6A). The impact of cellular oxidative stress following Pro treatment was then assessed using an assay to quantify malondialdehyde (MDA) levels. MDA is a by-product of lipid peroxidation that reflects accumulated cellular oxidative damage. MDA levels were significantly increased in *aox1a.aox1d* lines but not WT following 24 h of Pro treatment (Fig 6B). These results suggest a potential role for both AOX1a and AOX1d to maintain redox balance and limit oxidative stress during periods of rapid Pro catabolism.

**Figure 6:**
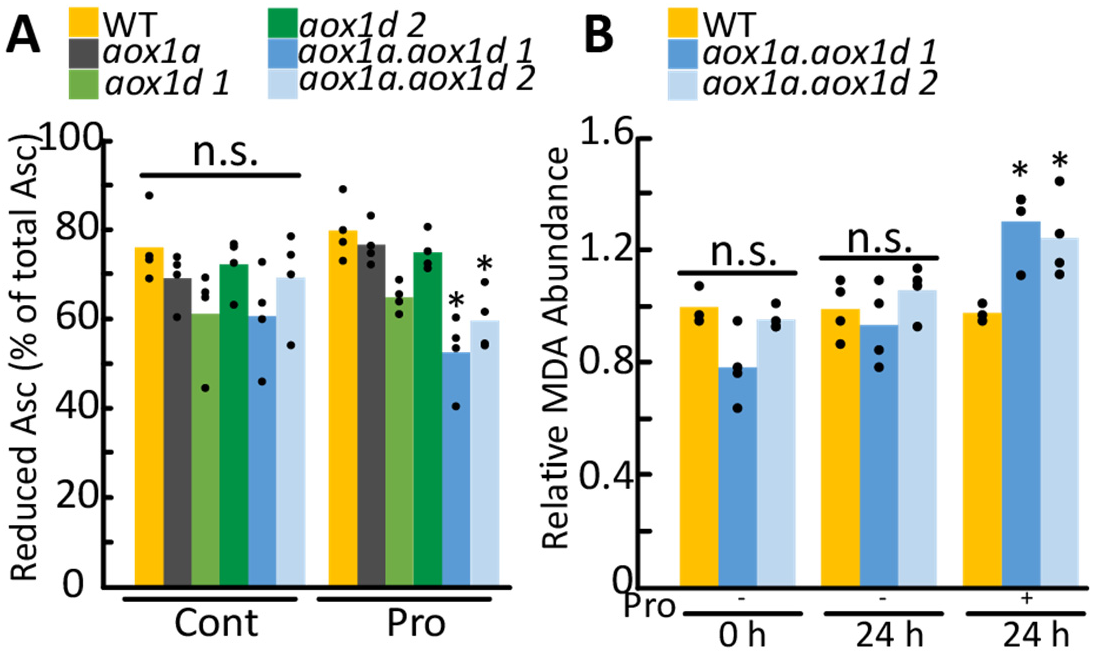
Quantification of ascorbate redox poise and MDA abundance following Pro exposure. **(A)** Redox status of the Asc pool in leaf discs in the presence and absence of Pro treatment. Replicate data points are shown (*n* = 4 independent biological replicates). Asterisks indicate significant treatment specific difference compared to WT (two-way ANOVA, p<0.05). **(B)** MDA abundance following exposure to Pro. Bars represent average MDA abundance relative to WT 0 h. Replicate data points are shown (*n* = 4 independent biological replicates). Asterisks indicate significant treatment specific difference compared to WT (two-way ANOVA, p<0.05).

### AOX1a and AOX1d aid in post-salinity recovery

To test the impact of loss of AOX1a and AOX1d at a physiological level, we assessed the ability of plants to recover from acute salt stress. Pro accumulates greatly in plant tissues during salinity stress and is quickly catabolised during subsequent recovery (Verslues and Sharma, 2010). Here, we show that Pro levels rose and fell in WT, *aox1a, aox1d*, and *aox1a.aox1d* leaves following seven days of salinity stress treatment and three days of recovery (Fig 7A and Supplemental Fig 4). The salinity stress treatment induced *AOX1a* and *AOX1d* expression in WT, similar to results from earlier studies (Smith et al., 2009, Van Aken et al., 2009) (Fig 7B). Much evidence suggests that AOX activity plays a key role in maintaining photosynthetic rates through cellular redox balancing (Vishwakarma et al., 2014, Vishwakarma et al., 2015) and photosynthetic electron transport rate (ETR) is known to be sensitive to saline stress (Stepien and Johnson, 2008). Therefore, we measured photosynthetic ETR during and after salinity stress. Following seven days of salinity stress, all genotypes showed similar reductions in their ETR response curves (Fig 7C). However, following salt removal and rewatering of the plants, both *aox1a.aox1d* lines displayed slower rates of recovery in their ETR response curves in comparison to WT or the *aox* single knockout lines (Fig 7C). These results lead us to conclude that *AOX1a* and *AOX1d* expression are responsive to conditions of elevated Pro *in vivo* and appear to be involved in optimising photosynthetic capacity during Pro drawdown and recovery from salt stress.

**Figure 7:**
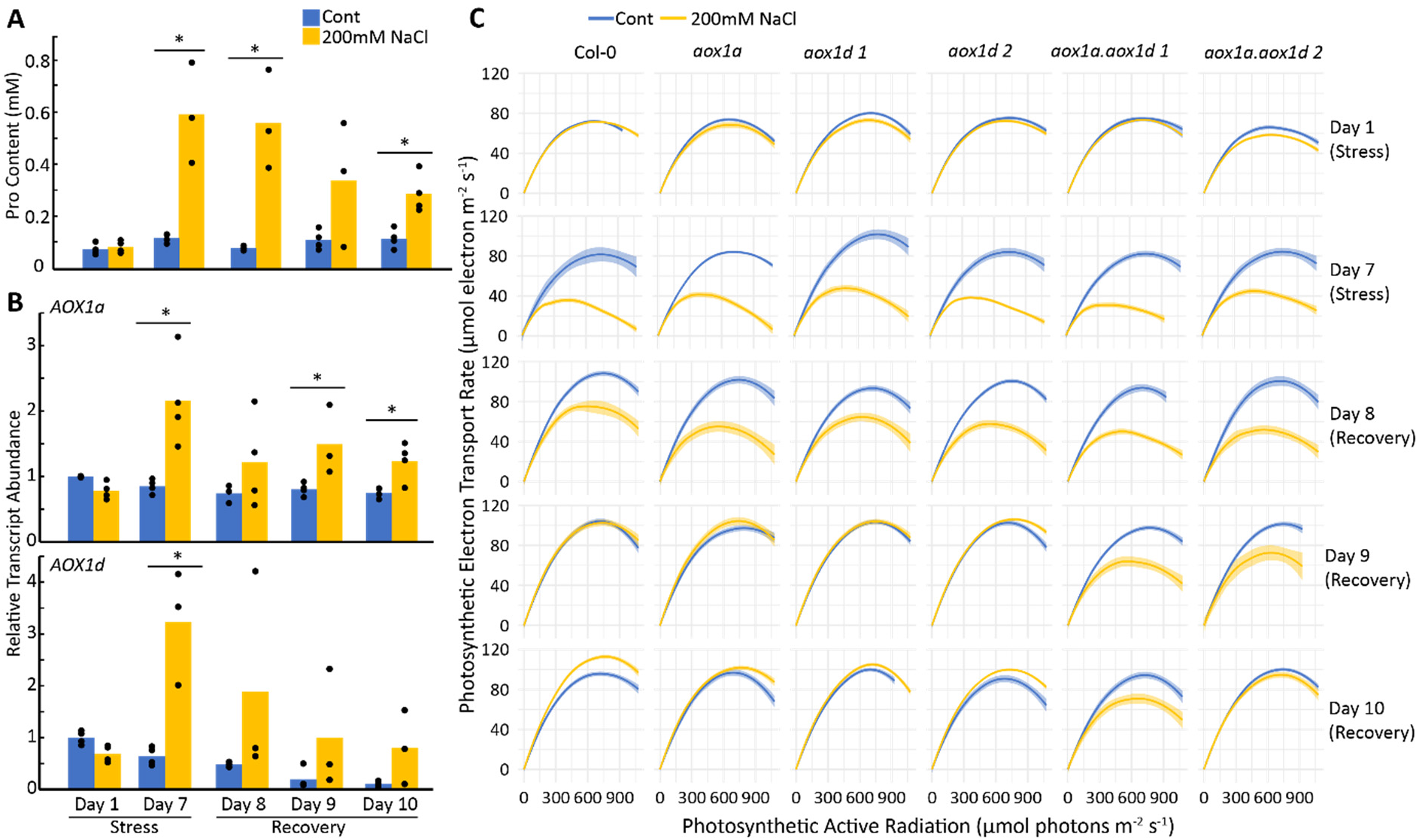
The effects of salinity and post-salinity recovery on Pro content, *AOX1a* and *AOX1d* expression levels and photosynthetic capacity. **(A)** Pro content from WT leaf discs obtained during salinity stress and post-salinity recovery. Bars represent average Pro content; replicate data points are shown (*n* ≥ 3). Asterisks represent significant differences between treatments (*t*-test; p < 0.05). **(B)** Transcript analysis of *AOX1a* and *AOX1d* from WT leaf discs obtained during salinity stress and post-salinity recovery. Bars represent average relative transcript abundance; replicate data points are shown (*n* ≥ 3). Asterisks represent significant differences between treatments (*t*-test; p < 0.05). **(C)** Photosynthetic ETR response curves of WT and *aox* lines during salinity stress and post-salinity recovery. Each curve represents the average Photosynthetic ETR response curve from four biological replicates with each biological replicate measured over six – 12 areas of interest on leaves. The shaded area shows the 95% confidence intervals.

## DISCUSSION

### AOX upregulation alleviates oxidative stress caused by Pro catabolism

This study explored the relationship between Pro catabolism and AOX activity in plants. It is well known that Pro accumulation in plant cells can lead to the upregulation of Pro catabolic enzymes (PDH and P5CDH), whose activities supply reductant directly or indirectly to the plant mETC (Cabassa-Hourton et al., 2016, Launay et al., 2019, O’Leary et al., 2019b). However, results from this and previous studies suggest that Pro exposure influences expression of other mETC components as well, in particular AOX (Fig 1B and Supplemental Table 1). Previously, mitochondria isolated from Pro-treated Arabidopsis seedlings displayed increased AOX capacity, which was particularly evident when Pro was used as the respiratory substrate (Cabassa-Hourton et al., 2016). Similarly, mitochondria isolated from senescing Arabidopsis leaves, which have high levels of ProDH expression, show a pronounced upregulation of Pro respiratory capacity *via* both COX and AOX pathways (Launay et al., 2019). Thirdly, we previously observed strong, Pro-dependent, respiratory stimulation of mature Arabidopsis leaf OCR that was inhibited by absence of both *ProDH1* and *ProDH2* (O’Leary et al., 2019b). Our results here confirm that these high rates of Pro catabolism also coincide with induced AOX capacity, thereby linking Pro exposure with AOX expression across several tissue types in Arabidopsis.

We initially hypothesised that enhanced AOX capacity may function to catalyse the increased electron flux emanating from Pro catabolism (Supplemental Fig 3B). We reasoned that although enhanced Pro catabolism could drive an approximate doubling of the respiration rate, there was no obvious need for a doubling of cellular ATP production; uncoupling the mETC could alleviate this imbalance. Previously, mitochondria isolated from Pro treated seedlings displayed a significant decrease in their ADP/O ratio, indicative of uncoupling (Cabassa-Hourton et al., 2016). However, here, leaves lacking detectable AOX expression still displayed full respiratory induction by Pro (Fig 3B and 4C). Therefore, while COX pathway capacity can be sufficient to catalyse the observed high rates of Pro catabolism in mature leaves (Fig 4C), AOX induction in WT was likely to contribute to Pro oxidation *in vivo*, for a reason beyond simply supplying additional respiratory rate.

The expression of AOX, specifically AOX1a, inversely correlates with both ROS abundance and oxidative damage in plant cells under different stress conditions (Maxwell et al., 1999, Pastore et al., 2001, Giraud et al., 2008, Vishwakarma et al., 2015, Gong et al., 2020). Since Pro catabolism *via* ProDH stimulates ROS production across eukaryotes (Donald et al., 2001, Zarse et al., 2012), we tested whether loss of AOX led to enhanced oxidative stress during Pro catabolism. Asc is an abundant antioxidant in plants and total Asc levels were strongly upregulated following Pro treatment in WT and AOX deficient plants, suggesting the induction of oxidative stress defence mechanisms (Fig 5 & Supplemental Table 2). However, the redox poise of the Asc:DHA pairing in *aox1a.aox1d* lines was significantly more oxidised, suggesting that AOX depleted tissues were more likely to be experiencing oxidative stress (Fig 6A). Indeed, more lipid-based oxidative damage was detected following prolonged Pro exposure in *aox1a.aox1d* lines compared to WT (Fig 6B). Therefore, enhanced AOX expression following Pro exposure is beneficial because it lowers oxidative stress associated with Pro catabolism.

#### Both AOX1a and AOX1d are responsible for AOX capacity in mETC upon Pro exposure

It is largely unknown if AOX isoforms other than AOX1a have specific purposes to fulfil in the respiratory pathways of plants. Do these AOX isoforms share the same functionality as *AOX1a* under certain perturbed metabolic conditions or do they act independently? While the upregulation of *AOX1d* transcripts during senescence (Clifton et al., 2006) and various stressful treatments (Strodtkotter et al., 2009, Feng et al., 2013, Demircan et al., 2020) is known, the perceived low abundance of AOX1d, compared to AOX1a, has limited further investigation (Fig 1C). In this study, we provided genetic and biochemical evidence of AOX1d’s role in Pro catabolism and its apparent functional redundancy to AOX1a in this role. Upon Pro exposure the fold increase of AOX1d was substantially greater than that of AOX1a (Fig 1B-D), and Pro-exposed *aox1a* leaves still displayed a significant increase in AOX capacity compared to control *aox1a* leaves (Fig 4A). By contrast, *aox1a.aox1d* leaves displayed no detectable AOX protein or activity (Fig 2B – D and Fig 4B) but rather displayed signs of enhanced redox imbalance compared to single *aox1a* or *aox1d* lines following Pro exposure (Fig 6A). Therefore, *AOX1d* functions in support of Pro catabolism, but judging from *AOX1d’s* expression profile from transcriptomic analyses (Clifton et al., 2006, Winter et al., 2007, Van Aken et al., 2009) its role may be more general in nature, such as supporting mETC redox balancing under stress.

A lack of characterised knockout lines have hindered progress in understanding the AOX1d isoform function in plants. The data we present here coupled to the complex evolutionary relationship of the AOX gene family in plants and evidence that AOX1a-c exist in a separate phylogenetic clade from AOX1d (Costa et al., 2014), suggest more attention is due to this isoform and its homologs. Whether a Pro catabolising function is conserved across AOX1d orthologs remains to be tested and the role of AOX1d in different metabolic stress scenarios also needs to be assessed. Results from this study also warn about the over interpretation of the effects of non-specific oxidase inhibitors such as SHAM and nPG on whole cells or intact tissues with regards to AOX function, again focusing on the need to use genetic resources to explore the roles of different AOX isoforms.

#### AOX1a and AOX1d aid in post-salinity recovery

The inhibition of AOX (*via* chemical or genetic means) leads to reduced photosynthetic capacity under stressful growth conditions (Giraud et al., 2008, Strodtkotter et al., 2009, Vishwakarma et al., 2014, Vishwakarma et al., 2015, Dinakar et al., 2016, Watanabe et al., 2016). Previous studies demonstrating the link between AOX expression and photosynthetic performance have only worked with *aox1a* plants (Strodtkotter et al., 2009, Vishwakarma et al., 2014, Vishwakarma et al., 2015, Dinakar et al., 2016, Watanabe et al., 2016). Here, we show that *aox1a.aox1d* leaves displayed a greater reduction in photosynthetic capacity during recovery from salinity stress compared to WT, *aox1a* or *aox1d* (Fig 7C). This pattern is consistent with our other measures of oxidative stress (Fig 6), and indicates a level of isoform redundancy between AOX1a and AOX1d (Fig 4A–B) with regards to enhancing photosynthetic capacity.

Interestingly, the negative impact of severe salinity stress on photosynthetic capacity did not differ between any of the genotypes while the stress was ongoing, but only during recovery (Fig 7C). During recovery from salinity, Pro can be returned back to control levels in a matter of days (Verslues and Sharma, 2010) (Fig 7A). The benefit of increased AOX capacity towards photosynthesis therefore coincides with this period of rapid Pro metabolism. In actively photosynthesizing cells, AOX is thought to optimize photosynthetic rates by better allowing the mETC to safely oxidize excess reductant exported by the plastid (Dahal et al., 2015, Del-Saz et al., 2018, Alber and Vanlerberghe, 2021). Vishwakarma et al. (2014) first observed higher NADH/NAD^+^ and NADPH/NADP^+^ ratios in high light treated *aox1a* Arabidopsis plants. Later, it was reported that antimycin A-treated *aox1a* plants displayed higher NADH/NAD^+^ ratios (Vishwakarma et al., 2015). Both observations suggest that the presence of AOX in WT plants allows for faster recycling of NAD^+^ and NADP^+^ to ensure optimal photosynthetic capacity. During salinity recovery, mitochondria may be required to balance excess reducing power generated concurrently from photosynthesis and from Pro catabolism. As observed in our study, the importance of mETC activity and AOX capacity toward cellular redox balancing may heightened under such conditions.

### Advantage of AOX activity is different metabolic scenarios

While the earlier view that AOX activity is, an energetically wasteful process has largely been dispelled, replacing this view with evidence for the specific metabolic benefits of AOX action have been slow. Our results demonstrate two specific outcomes of AOX expression that are beneficial in metabolism from an energetic standpoint. First, AOX activity can limit oxidative damage to cellular macromolecules under demanding respiratory conditions imposed by Pro catabolism (Fig 6B). The ATP savings from a reduced rate of macromolecule turnover due to oxidative damage will need to be weighed against the loss of ATP production because of AOX activity to fully explain why this trade-off is beneficial. Secondly, AOX action supports recovery of photosynthetic rate and therefore directly increases the overall energy/carbon budget available to the cell under conditions of recovery from salinity (Fig 7C). These expanded energetic accounting considerations are consistent with the phenotypical observation that AOX expression is generally positive for plant performance (Vanlerberghe, 2013, Dahal et al., 2015, Del-Saz et al., 2018, Selinski et al., 2018b).

## MATERIALS & METHODS

### Plant material and growth conditions

Arabidopsis (*Arabidopsis thaliana*) accession Col-0 (N6000) was used as wild-type (WT). The following mutant lines were used: *aox1a* (At3g22370; SAIL_030_D08; Giraud et al. (2008)), *aox1d 1* (At1g32350; SALK_203986), *aox1d 2* (GABI_529_D11) and *pdh1-1.pdh2-1* (At3g30775 and At5g38710; (Funck et al., 2010, O’Leary et al., 2019b)). Two independent *aox1a aox1d* lines were constructed by crossing *aox1a* with the two *aox1d* lines (*aox1a.aox1d 1* and *aox1a.aox1d 2*) (Supplemental Fig 2A). Primers used for genotyping of *aox1d 1* and *aox1d 2* can be found in Supplemental Table 3.

Seeds were sown on a 3:1:1 potting soil: perlite: vermiculite supplemented with slow releasing fertiliser and covered with transparent cover until seedlings are established. Seedlings were then transplanted into individual pots with soil kept moist with a regular watering schedule. Plants were grown in a controlled growth chamber maintaining a short-day photoperiod of 8h light (21°C): 16h dark (18 °C) (2300 h – 0700 h light) with a photon flux of 100 μmol m^-2^s^-1^ and a relative humidity of 65 %. For respiratory measurements leaf discs samples (7 mm in diameter) were taken from mature leaves of 8 – 10 weeks old plants, which were in the vegetative stage. For treatments prior to crude mitochondrial extracts, 8 – 10 weeks old leaves were cut into strips (approximately 4-5 strips per leaf) and floated adaxial side up on respiration buffer in the presence or absence of Pro.

For salt stress treatments, eight biological replicates (four control, four salt-treated) of all six genotypes (*i.e*. WT, *aox1a, aox1d 1, aox1d 2, aox1a.aox1d 1* and *aox1a.aox1d 2*) were placed in two separate square trays. Each individual pot contained 50g of soil. To induce salinity conditions, salinity-treated plants were watered with 1L of 200mM NaCl solution and control-treated plants were watered with 1L of water. Solution in both trays were replaced every day during the treatment period. For post-salinity recovery, both trays were subjected to multiple cycles of washing which included placing pots on paper towels for an hour and transferring them to 1L of water for another hour. Arabidopsis plants were subjected for salinity stress (for seven days) and re-watered (for three days).

### Respiratory measurements

Leaf discs respiratory measurements were performed using the Q2 oxygen sensor (Astec Global) in sealed 850 μL capacity tubes at 21 °C by the method of (O’Leary et al., 2019b). Leaf discs were sampled 4 h into their night period and floated on the adaxial side up on 600 μL of leaf respiration buffer (50mM HEPES pH 6.6, 10mM MES, 2mM CaCl_2_) with additional metabolites or chemicals as described. All Q2 experiments sampled a minimum of six leaf discs from different pots. To obtain R_N_ respiratory traces, 2 h moving slopes of O_2_ depletion over time were calculated.

Leaf discs respiratory measurements using the Clark-type oxygen electrode (Hansatech) were conducted following 14 h of Pro exposure, with eight leaf discs in 1.5 mL of respiration buffer (50mM HEPES, 10mM MES, 2mM CaCl_2_ and pH 6.6). AOX capacity was measured as the difference in OCR between the addition of 2 mM KCN and 4mM SHAM.

Mitochondrial respiration rate measurements were performed using the Clark-type oxygen electrode (Hansatech) with 120 - 240 μg of enriched mitochondria in 1 mL of mitochondrial respiration buffer (0.3M Sucrose, 5 mM KH_2_PO_4_, 10 mM TES, 10 mM NaCl, 2 mM MgSO_4_, 0.1 % (w/v) BSA in MilliQ water, pH 7.2 at 25 °C (Jacoby et al., 2015). To quantify Pro-dependent respiration, 250 μM ADP and 6 mM Pro were used to activate the mETC. To quantify Complex I & II dependent respiration, 5mM Succinate, 1mM NADH and 250 μM ADP were used to activate the mETC. Electron capacity *via* the COX pathway was measured as oxygen consumption rates sensitive to KCN treatment. KCN (2mM) was used to inhibit the COX pathway and determine AOX capacity. nPG (0.1mM) was used to inhibit the AOX pathway and determine background respiratory activity.

### Transcript analysis

Leaf discs were subjected metabolite incubations as described above. Following 12 hrs of metabolite exposure, leaf samples were collected and snap frozen in liquid nitrogen. For transcript analysis of WT plants during salinity stress, leaf samples were harvested after obtaining photosynthetic ETR curves at the respective timepoints. In all cases, two leaf discs were combined to form one biological replicate and powdered using a mixer mill. RNA extraction, cDNA synthesis and qRT-PCR were performed in accordance to O’Leary et al. (2019b). Analysis of amplification curves were done using LinregPCR (Ruijter et al., 2009). Absolute N_0_ values of three technical replicates were averaged for each of the biological replicates (*n* = 4). Relative transcript abundance was obtained by normalising to two housekeeping genes (*AtACT2* and *AtUBQ10*). The two normalised values were then combined using geometric mean calculation. Primers used in qRT-PCR can be found in Supplemental Table 3.

### Quantification of proline content

Two leaf discs were harvested from two leaves from each pot and combined to form one biological replicate. Leaf samples were powdered using a mixer mill and measurement of Pro content was performed as detailed previously in O’Leary et al. (2019b). Samples were extracted once in 200 μL of 80 % ethanol at 80 °C for 20 mins and spun down at full speed for one minute. Resulting supernatant (100 μL) was added to 200 μL of reaction mix (1 % (w/v) ninhydrin, 60 % (v/v) acetic acid, and 20 % (v/v) ethanol). The samples were incubated at 95 °C for 20 mins, cooled on ice and transferred to a microtiter plate for measurement of *A_520_*. Pro concentrations were interpolated using a standard curve.

### Extraction of crude mitochondria

Enriched mitochondria extracts were isolated at 4 °C from 12 g of mature leaves (8 – 10 weeks old) following a 12 hr exposure to metabolites. Extraction of crude mitochondria was performed as detailed in Millar et al. (2007). Harvested leaves were ground using a pre-chilled mortar and pestle and 120 ml of extraction buffer (10 mM TES, 0.3 M Sucrose, 25 mM tetrasodiumpyrophosphate, 2 mM EDTA, 10 mM KH2PO4, 1 % (w/v) polyvinylpyrrolidone 40, 1 % (w/v) BSA, adjusted to pH 7.5 with 85 % phosphoric acid. 20 mM Ascorbic acid and 5 mM Cysteine was added before use). Ground tissue was filtered through two layers of Miracloth and centrifuged for five minutes at 2,500 *g* at 4 °C. The supernatant was transferred and centrifuged for 20 minutes at 18,000 *g* at 4 °C. The resulting pellet was washed twice with sucrose wash buffer (0.3 M sucrose, 10 mM TES, 7.4 mM KH_2_PO_4_, adjusted to pH 7.5 with NaOH). Washing of pellet was carried out twice by subjecting samples to the same low and high-speed centrifugation mentioned above. Enriched mitochondria pellet was resuspended in 250 μL of sucrose wash buffer. Enriched mitochondrial protein was quantified using Coomassie Plus (Bradford) Assay Kit (Thermo Fisher Scientific) with BSA as a standard.

### Immunoblotting

Enriched mitochondria pellets were extracted in SDS sample buffer (4 % (w/v) SDS, 20 % glycerol, pH 6.8 125 mM Tris-HCl, 1 μL of bromophenol blue and 50 mM DTT) containing 2 mM PMSF and cOmplete protease inhibitor cocktail (Roche) according to manufacturer’s instructions. Samples were ran on a 12 % precast SDS-PAGE gel (Bio-rad, catalogue no. 456046), transferred onto PVDF membrane and blocked in 2 % skim milk. Blot was probed at 4 °C with orbital shaking overnight with anti-AOX antibody (Agrisera, catalogue no. AS04054) at 1:2000 dilution or anti-VDAC antibody (Agrisera, catalogue no. AS07212) at 1:5000 dilution. Detection of the mitochondrial VDAC protein was used as a loading control. After secondary antibody (Anti-Rabbit, Sigma Aldrich, catalogue no. A0545) incubation, blot was exposed to ECL Western Blotting Substrate (Thermo Fisher Scientific, catalogue no. 32106) and imaged using an Amersham 680 Imager CCD camera (GE Life Sciences).

### Oxidative stress measurements

Lipid peroxidation was quantified using the thiobarbituric acid reactive substances (TBARS) assay as detailed previously in Hodges et al. (1999), Calzadilla et al. (2016) with modifications. Following 24 hrs of Pro exposure, leaf disc samples were collected and snap frozen in liquid nitrogen. Four biological replicates were collected; each biological replicate had eight leaf discs combined and powdered using a mixer mill. 5% (w/v) meta-phosphoric acid (750 μL) and of 2% (w/v) butyl hydroxytoluene (15 μL) were added to the powdered samples, vortexed and centrifuged for 20 minutes at 15,000 *g* at 4 °C. Resulting supernatant (600 μL) was added to a mixture of of 2% (w/v) butyl hydroxytoluene (40 μL), 1% (w/v) thiobarbituric acid (200 μL) and 25% (v/v) HCl (200 μL) and heated at 95 °C for 30 minutes with shaking at 450 rpm. 1-butanol (700 μL) was added and the resulting mixture was centrifuged for 2 minutes at 15,000 *g*. Resulting supernatant (200 μL) was used for quantifying *A_532_* and *A_600_* using a spectrophotometer. Absorbance of MDA was calculated as the difference between *A_532_* and *A_600_*.

Ascorbate measurements were performed based on the methods of Foyer et al. (1983), Queval and Noctor (2007) with modifications. Following 4 h of Pro exposure, leaf samples were collected and snap frozen in liquid nitrogen. Four biological replicates were collected; each biological replicate had 10 leaf discs combined and powdered using a mixer mill. Ice cold 1M HClO4 (700 μL) was added to each sample and subjected to centrifugation for 10 minutes at 16,000 *g* at 4°C. Resulting supernatant (500 μL) was transferred into a new tube containing 50 μL of 0.2M NaH_2_PO_4_ (pH 5.6). The extract was then neutralised with 5M K_2_CO_3_ to pH 5~6 and subjected to centrifugation 5 minutes at 16,000g at 4°C. To assay reduced ascorbate, triplicate aliquots of 40 μL of neutralised supernatant were added to plate wells containing 55 μL of water and 100 μL 0.2M NaH_2_PO_4_ (pH 5.6) and *A_265_* was recorded using the spectrophotometer. To assay total ascorbate, 150 μL of neutralised supernatant was added to a mixture of 210 μL 0.12M NaH_2_PO_4_ (pH 7.5) and 20 μL of 25mM DTT and incubated at room temperature for 30 minutes in dark. Total ascorbate was then measured in the same manner as reduced ascorbate. Oxidised ascorbate is the difference between total and reduced ascorbate.

### Protein and metabolite mass spectrometry

Targeted mass spectrometry via multiple reaction monitoring (MRM) was employed to quantify mETC protein abundance. Pre-chilled 100 % acetone (400 μL) was added to 100 μg of enriched mitochondria and incubated overnight at – 20 °C. The resulting pellets were washed with acetone thrice. Samples were alkylated, trypsin digested, desalted and cleaned as previously described by Petereit et al. (2020). Samples were loaded onto an AdvanceBio Peptide Map column (2.1 × 250 mm, 2.7-μm particle size; part number 651750-902, Agilent), using an Agilent 1290 Infinity II LC System coupled to an Agilent 6495 Triple Quadrupole MS as described previously (James et al., 2019).The list of peptide transitions used for MRM is provided in Supplemental Table 4. Peptide abundances from each sample were normalised against VDAC.

Metabolite abundance was measured using gas-chromatography and liquid chromatography mass spectrometry as previously detailed in O’Leary et al. (2017) and Le et al. (2021), respectively.

### Photosynthetic electron transport rate measurement

Chloroplastic ETR measurements were performed using an IMAGING-PAM *M-Series* (MAXI version) (Walz; Effeltrich, Germany). ETR response curves were obtained using an inbuilt script (settings: 2, corresponding to the following PAR: 0, 1, 11, 21, 36, 56, 81, 111, 146, 186, 231, 281, 336, 396, 461, 531, 611, 701, 801, 926, 1076 ^1^, 20 s interval between each measurement). The ETR of technical replicates (six – 12 areas of interest) were combined to get the leaf ETR, and the ETR of different leaves for the same treatment were plotted to get the treatment ETR trend. A loess regression was adjusted to each curve with a 95% confidence interval using the R package ggplot2.

## Data availability

All data are provided in supplemental files

## Acknowledgements

We thank Ricarda Fenske for optimising the method and running the targeted MRM for the mitochondrial protein abundance measurements. We thank Chun-Pong Lee for a critical reading of the manuscript.

## Supplemental files

**Supplemental Figure 1:** Transcript abundance of *AOX1b, AOX1c* and *AOX2* after Pro exposure in WT leaf tissues.

**Supplemental Figure 2:** Characterisation of *aox1d* and *aox1a.aox1d 2* lines.

**Supplemental Figure 3:** The effect of SHAM and nPG on Pro-stimulated respiration

**Supplemental Figure 4**: The effect of salinity and post-salinity recovery on Pro content.

**Supplemental Table 1:** Relative abundance of mitochondrial proteins measured using targeted MRM.

**Supplemental Table 2:** Quantification of cellular metabolites in leaf discs following Pro treatment

**Supplemental Table 3:** Primers used for transcript analysis as outlined in the methods section.

**Supplemental Table 4:** The MRM transitions used to measure relative abundance of mitochondrial proteins

